# Efficient recovery of transgene-free edited tobacco using the transgene self‑elimination system via *de novo* regeneration

**DOI:** 10.64898/2026.10.24.761527

**Authors:** Jianduo Zhang, Weiran Li, Jing Lv, Xinruitong Liu, Luoping Wang, Yingying Qian, Yuying Liu, Feiyan Qi, Haoyi Xue, Xin Liu, Lele Deng, Fengjiao Wang, Jicang Xu, Kaili Xue, Xuepeng Liu, Qi Wu, Haiying Xiang, Changbo Dai

**Affiliations:** Raw Material Center of China Tobacco Yunnan Industrial Co., Ltd., Kunming, 650201, China; Tobacco Research Institute, Chinese Academy of Agricultural Sciences, Qingdao, 266101, China; Yunnan Academy of Tobacco Sciences, Kunming, 650032, China

**Keywords:** Genome editing, Transgene-free, Transgene Killer CRISPR, Agrobacterium-mediated regeneration, Tobacco

## Abstract

Elimination of integrated editing constructs remains a major challenge for generating transgene-free progeny in plant genome editing based on Agrobacterium-mediated regeneration. Here, we adapted the Transgene Killer CRISPR (TKC) concept to tobacco and established the transgene self-elimination system, in which CRISPR/Cas9 is coupled with developmentally regulated BARNASE cassettes. Three pU3TR-derived vectors carrying *DLL*-, *OsRP3*- or *PHAS*-driven *Barnase* regenerated normally after leaf-disc transformation. The DLL construct pEB138 showed the highest T_0_ editing frequency, with 56 of 69 independent T_0_ plants (81.16%) carrying mutations at one or both *NtDFR* target loci. Under a stringent lineage-level screen in which up to 48 T_1_ progeny were tested per T_0_ line, 36 of 43 pEB138 lineages (83.72%) yielded uniformly marker-negative progeny for the three assayed T-DNA markers, compared with 0 of 7 lineages transformed with the parental pU3TR vector. The platform also supported recovery of *dfr* mutants with altered flower pigmentation and *NtA622*-edited lines with reduced nicotine accumulation. Moreover, introduction of *DLL-iBarnase* into the independent pDC45 editing backbone did not impair shoot regeneration, and representative *NtBBL* progeny showed markedly reduced nicotine content together with loss of the selectable cassette. These results demonstrate that BARNASE-assisted transgene elimination can be integrated with Agrobacterium-mediated *de novo* regeneration in a dicot, enabling one-step recovery of transgene-free edited tobacco progeny.

---

For most crops, genome editing relies on *A*grobacterium-mediated transformation followed by *de novo* regeneration, a route that leaves the editing cassette stably integrated until it is removed by segregation in later generations. Edited plants that retain the CRISPR/Cas9 cassette blur the distinction between inherited and newly induced mutations, prolong nuclease exposure and complicate both breeding and regulatory assessment (Zhang et al., 2018). A single heterozygous T-DNA locus yields only about 25% transgene-free progeny after selfing, and every additional insertion further reduces this fraction. The Transgene Killer CRISPR (TKC) system overcame this limitation in rice by coupling the editing cassette to two suicide modules, 35S-driven CMS2 and REG2-driven BARNASE, which eliminate transgene-carrying male gametophytes and embryos, respectively (He et al., 2018). This principle was subsequently applied to multiplex gene-family editing (Yang et al., 2022), and was further refined in TKC2, which couples a RUBY reporter with the suicide modules to identify occasional escapees (Zhu et al., 2025). He et al. (2018) proposed that the strategy should be transferable to species transformed through tissue culture, although it was unlikely to suit the *Arabidopsis* floral-dip route. Other transgene-free strategies in *Arabidopsis* have instead relied on gamete-mediated transgene elimination introduced by floral dip (Cheng and Sun, 2025). Given the profound differences between monocots and dicots in gametophyte and embryo development (Zhao et al., 2017), whether a transgene self-elimination module can be coupled to routine *Agrobacterium*-mediated *de novo* regeneration in a dicot has remained an important practical question.

Previously, a CRISPR/Cas9-based mutant library was created in tobacco (*Nicotiana tabacum*), a model solanaceous species and a key plant chassis for plant synthetic biology that is routinely transformed by Agrobacterium-mediated de novo regeneration (Chen et al., 2023). However, fewer than 10% of T_1_ progeny were transgene-free, and most edited lines retained the integrated Cas9 construct, hampering genetic analysis and downstream breeding. This low recovery of transgene-free progeny is a key bottleneck, prompting us to develop an efficient transgene self-elimination strategy. Gamete- or seed-preferential promoters offer an attractive route for precise transgene self-elimination in reproductive tissues, but even residual expression during transformation and regeneration can abolish recovery of viable plantlets, the principal obstacle to transferring this approach to *de novo* regeneration-based systems. In previous work, we developed an optimized CRISPR/Cas9 system, pOREU3TR (designated pU3TR), which harbors an AtUB10-ROS1 visual marker to facilitate the screening of transgene-free plants (Zhang et al., 2023), but this strategy remains time-consuming and labor-intensive. Here, we upgraded the pU3TR to a transgene self-elimination system, achieving up to 83.72% marker-negative lineages in the T_1_ generation, approximately eight-fold higher than the transgene-free recovery previously achieved with the parental pU3TR vector, and rapidly generated transgene-free, low-nicotine tobacco lines.

To construct the transgene self-elimination system, we ultimately incorporated three optimal candidates (*DLL, OsRP3* and *PHAS*) through systematic analysis and functional screening of multiple putative reproductive-preferential promoters (Figure 1a and Table S1). Three derivative constructs, pEB138, pEB139 and pEB140, were generated to drive the expression of modified *BARNASE* with two introns (*iBarnase*) under the independent control of the *DLL, OsRP3* and *PHAS* promoters, respectively (Figure 1a and Figure S1). To validate transgene self-elimination systems, we selected tobacco *dihydroflavonol 4-reductase* (*NtDFR*) genes as editing targets. Loss-of-function mutants in *NtDFR* produce white flowers and enable straightforward phenotypic screening (Jiang et al., 2023). All four constructs, including the parental pU3TR vector, produced regenerating shoots after transformation with *Agrobacterium tumefaciens* (Figure 1b). Compared with the parental pU3TR control, which produced an average of 3.89 purple buds per explant, the *iBarnase-*bearing derivative constructs pEB138, pEB139 and pEB140 yielded comparable bud counts of 3.78, 4.00 and 3.11 per explant, respectively, indicating that self-elimination modules caused no measurable impairment of regeneration capacity under the tested conditions (Figure 1c). Target-locus genotyping of T_0_ regenerants revealed robust editing by pEB138 at the two *NtDFR* loci, *NtDFRa* (*Ntab0804470*) and *NtDFRb* (*Ntab0921540*), the subgenome copies of *NtDFR* in the allotetraploid tobacco genome. Of 69 independent pEB138-derived T_0_ plants, 45 (65.22%) and 50 (72.46%) carried targeted mutations at these two loci, respectively, with 22 (31.88%) and 26 (37.68%) classified as homozygous or biallelic (Ho/Bi) mutations. When all mutation types were pooled across the two loci, 56 of 69 plants contained targeted edits, an overall editing efficiency of 81.16%, which was comparable with the 80.95% achieved by the parental pU3TR control. For pEB139, 8 of 21 regenerants (38.10%) and 7 of 21 regenerants (33.33%) carried edits at the two loci, with no Ho/Bi mutations recovered. In the pEB140 group, the corresponding editing frequencies were 7 of 18 (38.89%) and 10 of 18 (55.56%), also without Ho/Bi events (Figure 1d). Thus, pEB138 maintained uncompromised regeneration capacity with high editing performance, and was therefore selected for detailed progeny-based analysis.

**Figure 1.**
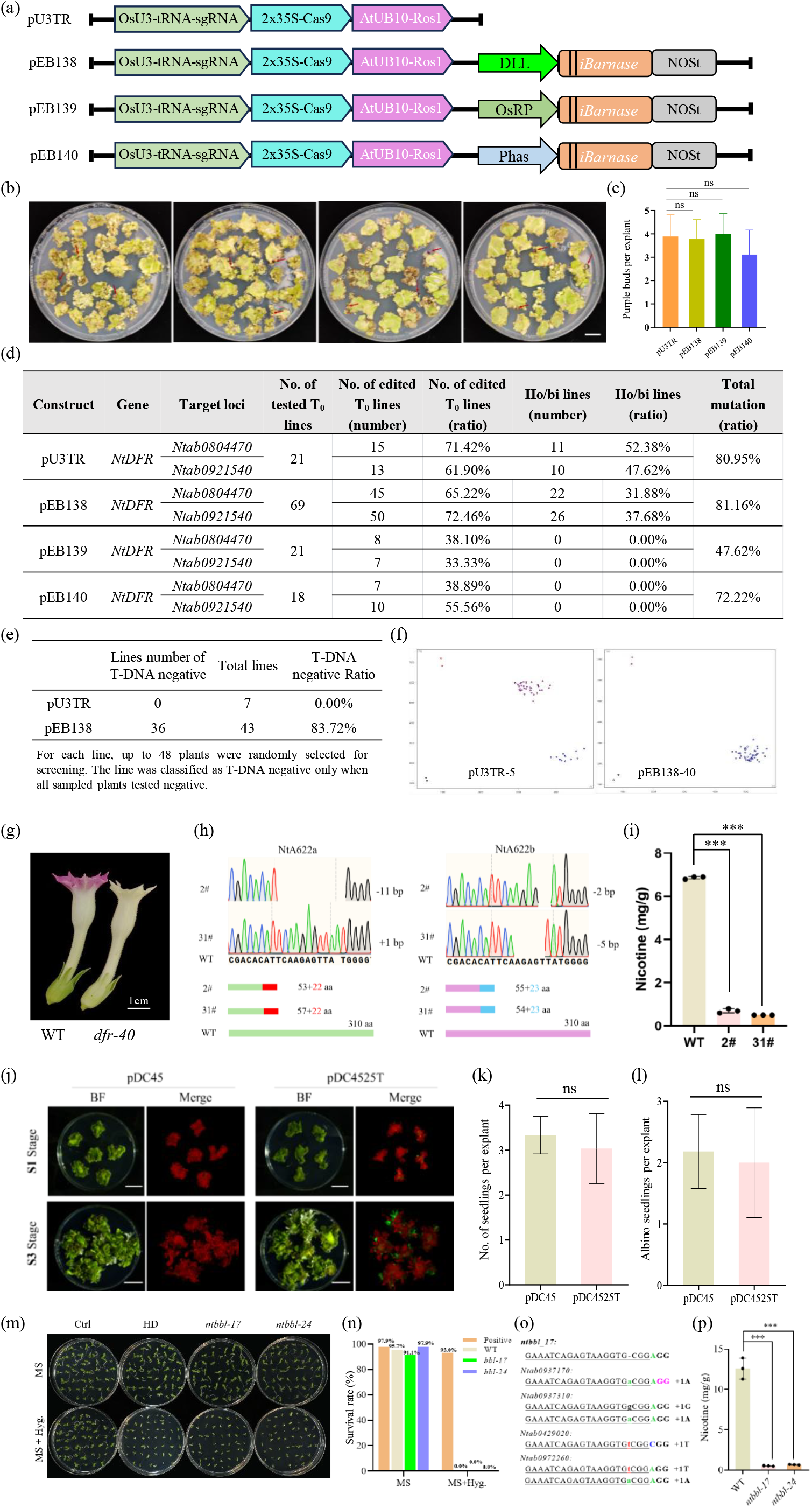
A transgene self-elimination system enables one-step recovery of transgene-free genome-edited tobacco via Agrobacterium-mediated *de novo* regeneration. (a)Schematic representation of the parental CRISPR/Cas9 vector pU3TR and three derivative constructs, pEB138, pEB139, and pEB140, carrying an intron-containing *iBarnase* expression cassette driven by the DLL, OsRP3, and PHAS promoters, respectively. (b)Representative shoot regeneration images from tobacco leaf explants transformed with pU3TR, pEB138, pEB139, or pEB140 vectors. Red arrows indicate regenerated purple shoots. (c)Quantification of purple buds regenerated per explant using different constructs. Data are presented as the mean ± standard deviation (SD) from at least three independent replicates. Statistical significance between four constructs was determined using Student’s t-test, with ns indicating not significant. The same statistical analysis was applied below. (d)Genome-editing efficiencies of the four constructs targeting two *NtDFR* loci (Ntab0804470 and Ntab0921540) in T_0_ tobacco plants. Numbers and percentages of edited plants and homozygous/biallelic (Ho/Bi) mutants are shown for each target. The total mutation frequency represents the percentage of T_0_ plants carrying mutations at either or both target loci. (e)Recovery of T-DNA-negative T_1_ progeny from pU3TR-and pEB138-transformed tobacco. For eachindependent T_0_ line, up to 48 randomly selected T_1_ seedlings were screened for vector-associated markers. A line was classified as T-DNA-negative only when all sampled seedlings tested negative. (f)Representative KASP genotyping plots of T_1_ progeny derived from pU3TR-5 and pEB138-40. Each point represents an individual progeny sample. (g)Representative flower phenotypes of wild-type (WT) and *NtDFR*-edited (*dfr-40*) tobacco plants, showing loss of floral pigmentation in the edited mutant. Scale bar, 1 cm. (h)Representative Sanger sequencing chromatograms showing mutations at the NtA622a and NtA622b loci in two independently edited lines (2# and 31#). The corresponding nucleotide insertions or deletions and predicted truncated protein products are indicated. aa, amino acids. (i)Nicotine contents in WT tobacco and the NtA622-edited lines 2# and 31#. Data are presented as the mean ± SD with three independent replicates. Statistical significance between WT and NtA622-edited lines was determined using Student’s t-test. Significance levels are indicated as follows: ***P < 0.001. (j)Representative images of tobacco tissues transformed with pDC45 or its *DLL-iBarnase*-containing derivative pDC4525T during shoot regeneration at the S1 and S3 stages. (k)Quantification of seedlings regenerated per explant from pDC45-and pDC4525T-transformed explants. (l)Quantification of albino seedlings regenerated per explant from pDC45-and pDC4525T-transformed explants. (m)Seed germination of control (Ctrl), HD, *ntbbl-17* and *ntbbl-24* on MS medium with or without hygromycin. (n)Survival rates of the indicated genotypes on MS medium with or without hygromycin. (o)Representative target-site sequences showing insertions at multiple *NtBBL* loci. (n)Nicotine contents in WT tobacco and the *ntbbl* mutants *ntbbl-17* and *ntbbl-24*. Data are presented as the mean ± SD with three independent replicates. Statistical significance was determined using Student’s t-test, with ***P < 0.001.

To quantify recovery of transgene-free edited progeny, up to 48 T_1_ seedlings were sampled from each independent T_0_ lineage and genotyped by Kompetitive Allele-Specific PCR (KASP) for three vector-associated markers targeting the sgRNA expression unit, the CaMV 35S promoter and the NOS terminator, representing the 5′-terminal, middle and 3′-terminal regions of the T-DNA, respectively. A lineage was scored as T-DNA-negative only when all selected progeny tested negative for all assayed markers. Under this stringent transgene-free definition, 36 of 43 pEB138 lineages (83.72%) met the criterion, whereas none of the seven lineages carrying the parental pU3TR vector satisfied it (Figure 1e, f). These results reveal strong enrichment of transgene-free edited progeny by the *DLL-iBarnase* configuration. Nevertheless, 7 of 43 pEB138 lineages still retained the vector markers, indicating that elimination was not absolute. The residual markers most likely reflect position effects of T-DNA insertion or attenuated DLL promoter activity, leading to *Barnase* expression below the threshold required for complete elimination.

Beyond editing efficiency, the workflow yielded stable trait-associated phenotypes. At the *NtDFR* loci, recovered alleles included small insertions and deletions, many of which are predicted to cause frameshifts. Edited plants displayed a pronounced loss of floral pigmentation relative to wild type, consistent with disruption of anthocyanin biosynthesis (Figure 1g). The *NtA622* gene encodes a key enzyme catalyzing the condensation of the pyridine and pyrrolidine rings of nicotine, and its loss of function produces a low-nicotine phenotype (Chang et al., 2026). In an independent application, the nicotine-biosynthesis homologues *NtA622a* and *NtA622b* were edited, and representative lines 2# and 31# carried distinct frameshift mutations at the target sites (Figure 1h). Nicotine content was markedly lower in both edited lines than in wild-type plants (Figure 1i). These phenotypic and biochemical results show that the pEB138 self-elimination system enables one-step generation of low-nicotine, transgene-free tobacco lines.

We next tested whether the elimination module could be transferred to a different editing architecture. pDC45 is a previously reported high-efficiency dicot editing vector in which a calreticulin-associated promoter configuration enhances CRISPR/Cas activity during regeneration (Li et al., 2025). pDC4525T was generated from pDC45 by replacing the hygromycin resistance cassette with a hygromycin-P2A-GFP cassette and adding the *DLL-iBarnase* module (Figure S2). Tissues carrying either pDC45 or pDC4525T developed normally through the S1 and S3 stages, with reporter fluorescence retained in pDC4525T-transformed explants during shoot development (Figure 1j), and the numbers of regenerated and albino seedlings per explant were comparable between the two constructs (Figure 1k, l), further indicating that the *Barnase* cassette did not impair regeneration. We then used pDC4525T to edit tobacco *berberine bridge enzyme-like* (*NtBBL*) genes, which encode the enzymes determining the stereochemistry of nicotine. Progeny from edited lines germinated on non-selective medium but were sensitive to hygromycin (Figure 1m, n). Representative target-site sequencing confirmed insertions at multiple *NtBBL* loci (Figure 1o), and nicotine content was markedly reduced in the *ntbbl* mutants (Figure 1p). These results are consistent with loss of the selectable editing cassette in the tested progeny.

In this study, we identified a T-DNA self-elimination element that is well suited to tobacco, coupling robust target-site editing with efficient germline removal of the editing construct and thereby enabling one-step recovery of transgene-free edited progeny. Although *Arabidopsis* is also a dicot, its floral-dip transformation route differs fundamentally from the regeneration-based transformation used here for tobacco, so strategies relying on passive transgene segregation cannot be directly applied to our workflow. The present system accordingly differs from the gamete-mediated transgene-free strategy reported in Arabidopsis via floral dip (Cheng and Sun, 2025), and from DNA-free ribonucleoprotein delivery, which avoids T-DNA integration but can be limited by delivery and regeneration requirements (Woo et al., 2015). A particular strength of the transgene self-elimination system is its simplicity, with a single DLL-driven BARNASE module that simultaneously eliminates transgene-carrying gametes and embryos, thereby simplifying the architecture of the rice TKC system, which relies on two separate suicide cassettes (He et al., 2022; Cheng and Sun, 2025). This extends the classic principle of tissue-preferential cytotoxic-ribonuclease expression for gamete ablation (Mariani et al., 1990) to the efficient enrichment of transgene-free edited progeny (He et al., 2022). Under the stringent criterion of up to 48 genotyped progeny per lineage, 36 of 43 (83.72%) pEB138 lineages yielded uniformly marker-negative T_1_ progeny, corresponding to 87.88% (1754/1996) of genotyped seedlings, whereas none of the seven pU3TR lineages did so. The design is also readily portable, since replacing DLL with species-specific reproductive- or seed-preferential promoters should adapt the module to any crop regenerated via Agrobacterium-mediated transformation, a route shared by many crops such as tomato, potato and pepper. Finally, the system complements our previously optimized pU3TR and pDC45 backbones by overcoming low T-DNA-negative recovery and rapidly delivering low-nicotine tobacco lines as proof-of-concept products. This study establishes the first single-expression-unit transgene self-elimination system, enabling efficient recovery of transgene-free edited progeny via Agrobacterium-mediated *de novo* shoot regeneration in a dicot.

## Supporting information

Suppemental figure

Suppemental table

## Acknowledgments

This study was supported by grants from Yunnan Daguan Laboratory (No. YNDG202401YZ01) and the Agricultural Science and Technology Innovation Program (ASTIP-TRIC02, CAAS-TRIC-2026-01).

## Author contributions

J.Z. and C.D. conceived and designed the study. J.Z. performed most of the experimental work, with the other authors performing part of the experiments and assisting with data analysis. C.D. and Q.W. wrote the manuscript. All authors read and approved the final version.

## Conflict of interest

The authors declare no conflict of interest.

## Data availability statement

All data supporting the findings of this study are contained within the article and its supplementary material.

## Notes

### Competing Interest Statement

The authors have declared no competing interest.

