## Supplemental methods for "Efficient recovery of transgene-free edited tobacco using the transgene self‑elimination system via *de novo* regeneration"

### *Plant materials and growth conditions*

Wild-type tobacco (*Nicotiana tabacum* cv. Honghuadajinyuan, hereafter HD) was used as the transformation recipient. Seeds were surface-sterilized with 75% ethanol for 30 s and 5% sodium chlorite for 8 min, rinsed four times with sterile water, and sown on Murashige and Skoog (MS) basal medium (pH 5.8) containing 30 g/L sucrose and 7 g/L agar. Seedlings were grown at 25°C under a 16 h light/8 h dark photoperiod. $T_0$ regenerated plants and $T_1$ progeny were grown in a greenhouse under the same temperature and photoperiod conditions.

### *Vector construction*

The basic editing vector pU3TR was modified from the pOREU3TR backbone and harbors an OsU3-tRNA-sgRNA expression cassette as well as an AtUB10-ROS1 visual reporter marker, as previously described (Zhang et al., 2023). To generate transgene self-elimination vectors, three complete expression cassettes were commercially synthesized. Each synthetic cassette consisted of one putative reproductive- or seed-specific promoter, a modified BARNASE sequence containing two introns (*iBarnase*), and a NOS terminator. The full-length cassettes were independently cloned between the *KpnI* and *BsrGI* restriction sites of the pU3TR backbone to generate three derivative vectors pEB138, pEB139 and pEB140, in which BARNASE expression was driven by the *DLL* (Polóniová et al., 2015), *OsRP3* (Zhou and Fan, 1993), and *PHAS* (van der Geest et al., 1996) promoters, respectively.

Vector pDC4525T was constructed based on the pDC45 backbone plasmid (Li et al., 2025). A synthetic hygromycin-P2A-mGFP expression cassette was assembled to function as a dual selectable marker. This marker cassette enables transgenic screening by hygromycin antibiotic resistance and facilitates rapid visual identification of positive transformants through mGFP fluorescence detection. The resulting intermediate construct was designated pDC4525. For the transgene self-elimination module, the *DLL-iBarnase-tNOS* expression cassette derived from pEB138 was amplified and integrated into the *KpnI* restriction site of pDC4525 using homologous recombination methods.

An sgRNA targeting a conserved coding region of *NtDFR* (5′-GCTCCAGTGACGCAAAC-CGTCGG-3′) was designed and inserted into the pU3TR or TKC backbone. All assembled vectors were verified by restriction digestion and Sanger sequencing. All oligonucleotide primers designed for vector assembly and amplification are listed in Table S4.

### *Agrobacterium-mediated tobacco transformation*

The TKC vectors were each introduced into *Agrobacterium tumefaciens* EHA105 by the freeze–thaw method, and Agrobacterium-mediated leaf-disc transformation was performed following our established procedure with minor modification (Li et al., 2025; Zhang et al., 2023). In brief, an overnight culture was resuspended in MSO medium to $OD_{600}$ = 0.6–0.8 with 20 mg/L acetosyringone; 0.8–1.0 cm leaf explants from 3–4-week-old aseptic seedlings were infected for 10 min, blotted dry, and co-cultivated on MS + 1-2.5 mg/L 6-BA + 0.1-0.25 mg/L NAA for 2 days in the dark. Explants were then selected on the same medium supplemented with 5 mg/L hygromycin or 50 mg/L kanamycin and 250 mg/L Timentin and were subcultured every 12 days. Shoots (1–1.5 cm) were rooted on MS + 0.05 mg/L NAA + 10 mg/L hygromycin or 100 mg/L kanamycin, and rooted $T_0$ plantlets were acclimatized and transferred to greenhouse soil.

### *Detection of mutations in $T_0$ plants*

Genomic DNA was extracted from leaf tissue of $T_0$ regenerated plants using the CTAB method. The *NtDFR* target region was amplified by PCR using site-specific primers flanking the sgRNA target site. PCR products were analyzed by Sanger sequencing. Mutation types were classified as biallelic (Bi), homozygous (Ho), heterozygous (Het), or wild-type (WT) based on sequencing chromatograms or high-throughput sequencing. The total mutation rate was calculated as the proportion of $T_0$ plants carrying any mutation at the target site.

### *Screening of T-DNA-free $T_1$ plants*

$T_0$ plants were self-pollinated, and $T_1$ seeds were harvested. For each independent $T_0$ lineage, up to 48 $T_1$ plants were randomly selected for genotyping. Genomic DNA was extracted and screened by PCR using three independent T-DNA marker primer pairs: one pair amplifying the CaMV 35S promoter (P35S), one amplifying the KfgS internal marker, and one amplifying the

NOS terminator (tNOS). A lineage was classified as T-DNA-negative only when all tested plants showed no amplification for all three markers. The T-DNA-negative ratio was calculated as the number of T-DNA-negative lineages divided by the total number of lineages screened.

**High-throughput KASP genotyping for T-DNA detection**

To enable high-throughput screening of the large $T_1$ population, residual T-DNA was genotyped using Kompetitive Allele-Specific PCR (KASP) markers performed on a Douglas Scientific Array Tape platform. Approximately 1 $cm^2$ of leaf tissue was collected from each plant and put into 96-well plates, freeze-dried, and ground with a 4-mm steel bead at 1,400 rpm for 2 min. Genomic DNA was extracted using an automated magnetic-bead-based plant DNA extraction protocol: 400 μL of lysis buffer was added and incubated for 1h, centrifuged at 3,600 rpm for 10 min, and DNA was purified. DNA quality was qualified and only samples with a concentration >10 ng/μL and an A260/280 ratio of 1.8–2.2 were used.

A two-stage pooled-then-individual strategy was adopted. In the first (pooled) stage, DNA from four individual plants of the same line was bulked into one pool, and all pools were genotyped. Any pool in which at least one T-DNA marker showed positive signal was processed in a second (single-plant) stage in which each individual was re-genotyped independently.

Three T-DNA-specific KASP markers were designed using BatchPrimer3 against the CaMV 35S promoter (35S-3), the KfgS internal element (KfgS-1) and the NOS terminator (TNOS-4). Each T-DNA marker was run as a duplex with an endogenous tobacco reference marker (L25). The T-DNA-specific forward primer carried the FAM fluorescence adapter (5′-GAAGGTGACCAAGTTCATGCT-3′), whereas the L25-specific forward primer carried the HEX fluorescence adapter (5′-GAAGGTCGGAGTCAACGGATT-3′); each primer pair was completed by its common reverse primer (Table S4). The FAM channel therefore reported the presence of T-DNA, whereas the HEX (L25) channel served as an internal amplification control confirming DNA quality.

PCR reactions were assembled by a NEXAR liquid-handling module in a total volume of 0.8 μL containing 2× KASP Master Mix (1× final), 0.375 μM common primer, 0.15 μM of each tailed allele-specific primer, and 10–20 ng of dried template DNA. Touchdown PCR was performed on a SOELLEX thermal-cycling module (94 °C for 15 min; 10 touchdown cycles of 94 °C for 20 s and 61→55 °C, −0.6 °C per cycle, for 60 s; followed by 26 cycles of 94 °C for 20 s and 55 °C for 60 s; additional cycles were appended when cluster separation was incomplete). End-point fluorescence was scanned with an ARAYA reader, and genotype clusters were called with INTELLICS software. Empty-vector DNA and wild-type tobacco DNA were included on every array as the transgene-positive and transgene-negative controls, respectively.

For single-plant calls, an individual was classified as T-DNA-negative when at least two of the three markers were negative (and none gave a conflicting positive call), T-DNA-positive when at least two markers were positive, and undetermined (N) when at least two markers failed. A line was classified as completely T-DNA-free only when every single plant within that line was negative for all three T-DNA markers; the T-DNA-negative ratio of each line was then calculated as the number of completely T-DNA-free lines divided by the total number of lines screened. This KASP-based two-stage workflow was used in parallel with the conventional three-marker PCR screening described above and yielded concordant genotypes.

### *Sanger sequencing and mutation analysis*

For T-DNA-free $T_1$ plants, the *NtDFR* target region was amplified and subjected to Sanger sequencing. Insertion/deletion (Indel) mutations were identified by aligning sequencing reads against the wild-type reference sequence. Mutations causing frameshifts were predicted to result in loss of function of the NtDFR protein.

### *Phenotypic observation*

Flower pigmentation phenotypes of wild-type and T-DNA-free *NtDFR* mutant plants were observed and photographed at the reproductive growth stage in the greenhouse.

Fluorescence and bright-field imaging of tobacco regenerating explants at S1 and S3 stages was performed on a Leica M205 FA stereo fluorescence microscope (Leica Microsystems, Wetzlar, Germany) fitted with a TL5000 Ergo transmitted-light base and LMT260 motorized XY scanning stage. Overlapping tile images were automatically acquired and assembled into panoramic composite images using the tile-scan mosaic function within LAS-X software. Bright-field and fluorescence channels were captured separately and merged to generate overlay images.

### *Statistical analysis*

Data are presented as numbers and percentages or as the mean ± standard deviation (SD) from at least three independent experimental replicates. Statistical significance between the control and treatment groups was determined using Student's t-test. Significance levels are indicated as follows: *$P < 0.05$, **$P < 0.01$, and ***$P < 0.001$; ns, not significant.
