## Supplementary material for "Efficient recovery of transgene-free edited tobacco using the transgene self‑elimination system via *de novo* regeneration": Suppemental figure

### Slide 1
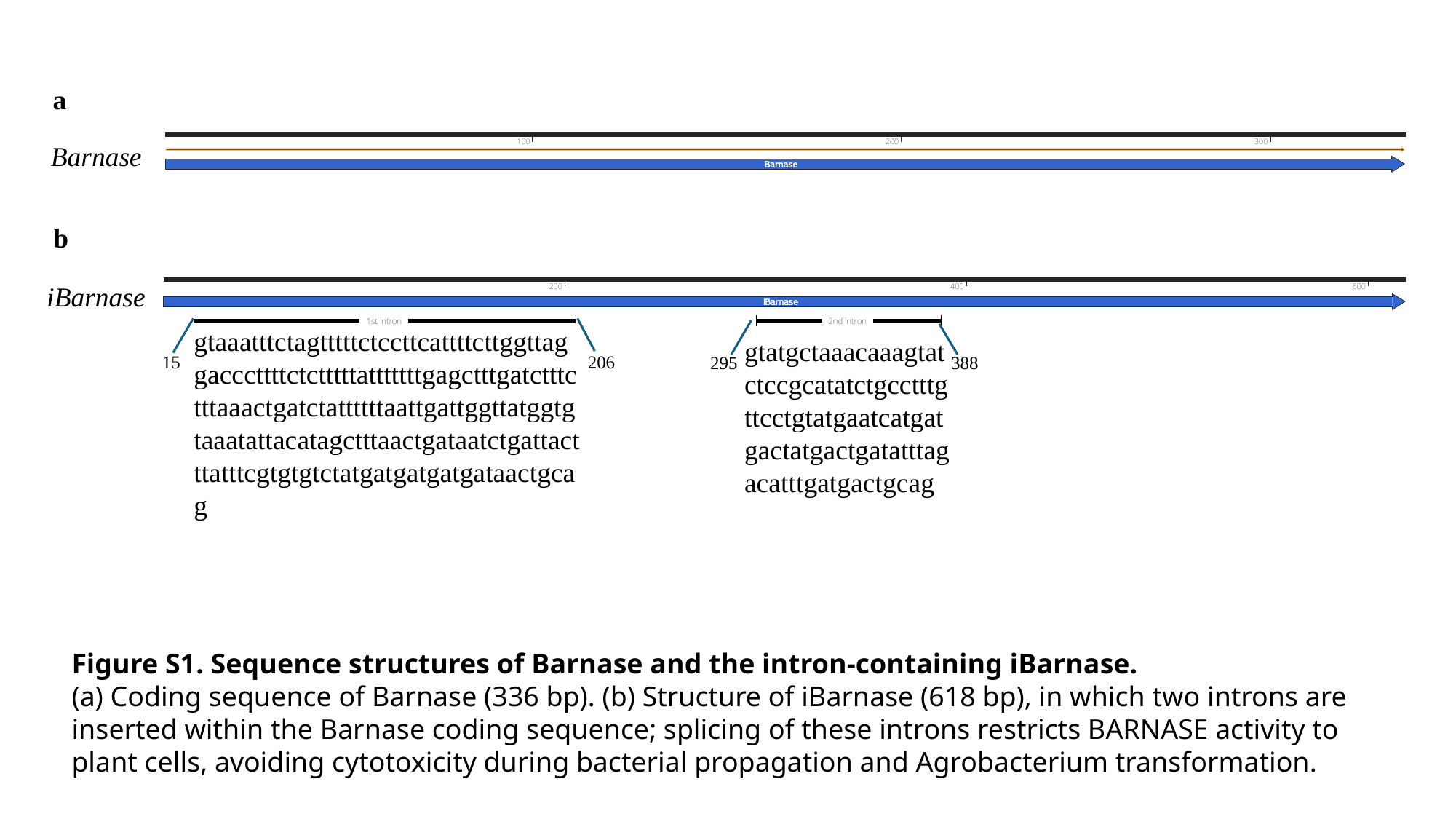

a
Barnase
b
iBarnase
gtaaatttctagtttttctccttcattttcttggttaggacccttttctctttttatttttttgagctttgatctttctttaaactgatctattttttaattgattggttatggtgtaaatattacatagctttaactgataatctgattactttatttcgtgtgtctatgatgatgatgataactgcag
gtatgctaaacaaagtatctccgcatatctgcctttgttcctgtatgaatcatgatgactatgactgatatttagacatttgatgactgcag
15
206
388
295
Figure S1. Sequence structures of Barnase and the intron-containing iBarnase.
(a) Coding sequence of Barnase (336 bp). (b) Structure of iBarnase (618 bp), in which two introns are inserted within the Barnase coding sequence; splicing of these introns restricts BARNASE activity to plant cells, avoiding cytotoxicity during bacterial propagation and Agrobacterium transformation.

### Slide 2
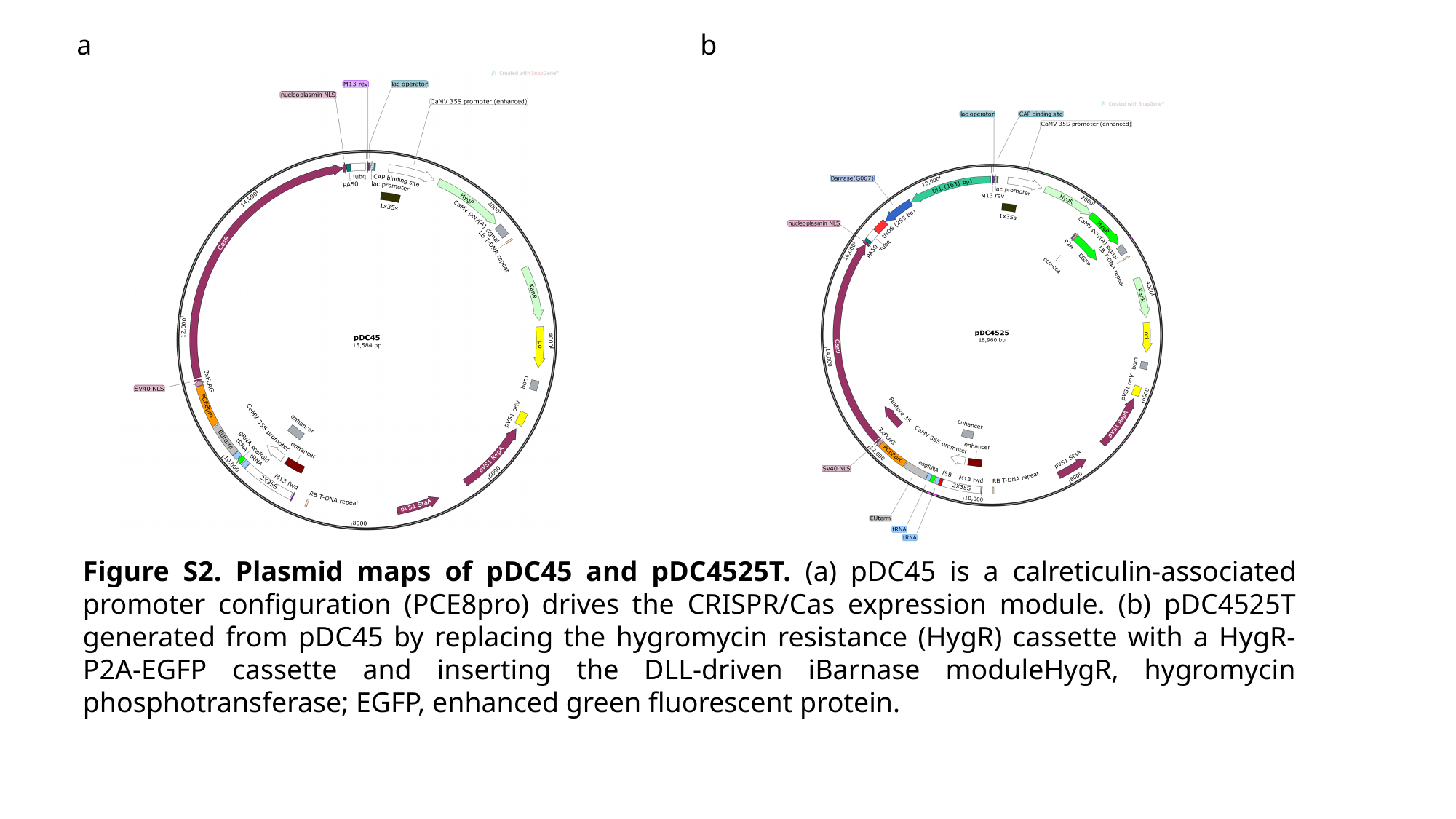

a
b
Figure S2. Plasmid maps of pDC45 and pDC4525T. (a) pDC45 is a calreticulin-associated promoter configuration (PCE8pro) drives the CRISPR/Cas expression module. (b) pDC4525T generated from pDC45 by replacing the hygromycin resistance (HygR) cassette with a HygR-P2A-EGFP cassette and inserting the DLL-driven iBarnase moduleHygR, hygromycin phosphotransferase; EGFP, enhanced green fluorescent protein.
